# Substrate profiling of the metalloproteinase ovastacin – Implications for its physiological function in mammalian fertilization

**DOI:** 10.1101/2022.12.06.519252

**Authors:** Matthias Felten, Ute Distler, Nele v. Wiegen, Mateusz Łącki, Christian Behl, Stefan Tenzer, Walter Stöcker, Hagen Körschgen

## Abstract

The metalloproteinase ovastacin is released by the mammalian egg upon fertilization and cleaves a distinct peptide bond in zona pellucida protein 2, a component of the enveloping extracellular matrix. This limited proteolysis causes zona pellucida hardening, abolishes sperm binding and thereby regulates fertility. Accordingly, this process is tightly controlled by the plasma protein fetuin-B, an endogenous competitive inhibitor. At present, little is known about how the cleavage characteristics of ovastacin differ from closely related proteases. Physiological implications of ovastacin beyond ZP2 cleavage are still obscure. In this study, we employed N-terminal amine isotopic labeling of substrates (N-TAILS) contained in the secretome of mouse embryonic fibroblasts to elucidate the substrate specificity and the precise cleavage site specificity. Furthermore, we were able to unravel the physicochemical properties governing enzyme-substrate interactions. Eventually, we identified several potential physiological substrates with significance for mammalian fertilization. These data suggest that ovastacin might regulate sperm-oocyte interaction and fertility beyond zona pellucida hardening.

**Graphical Abstract:** 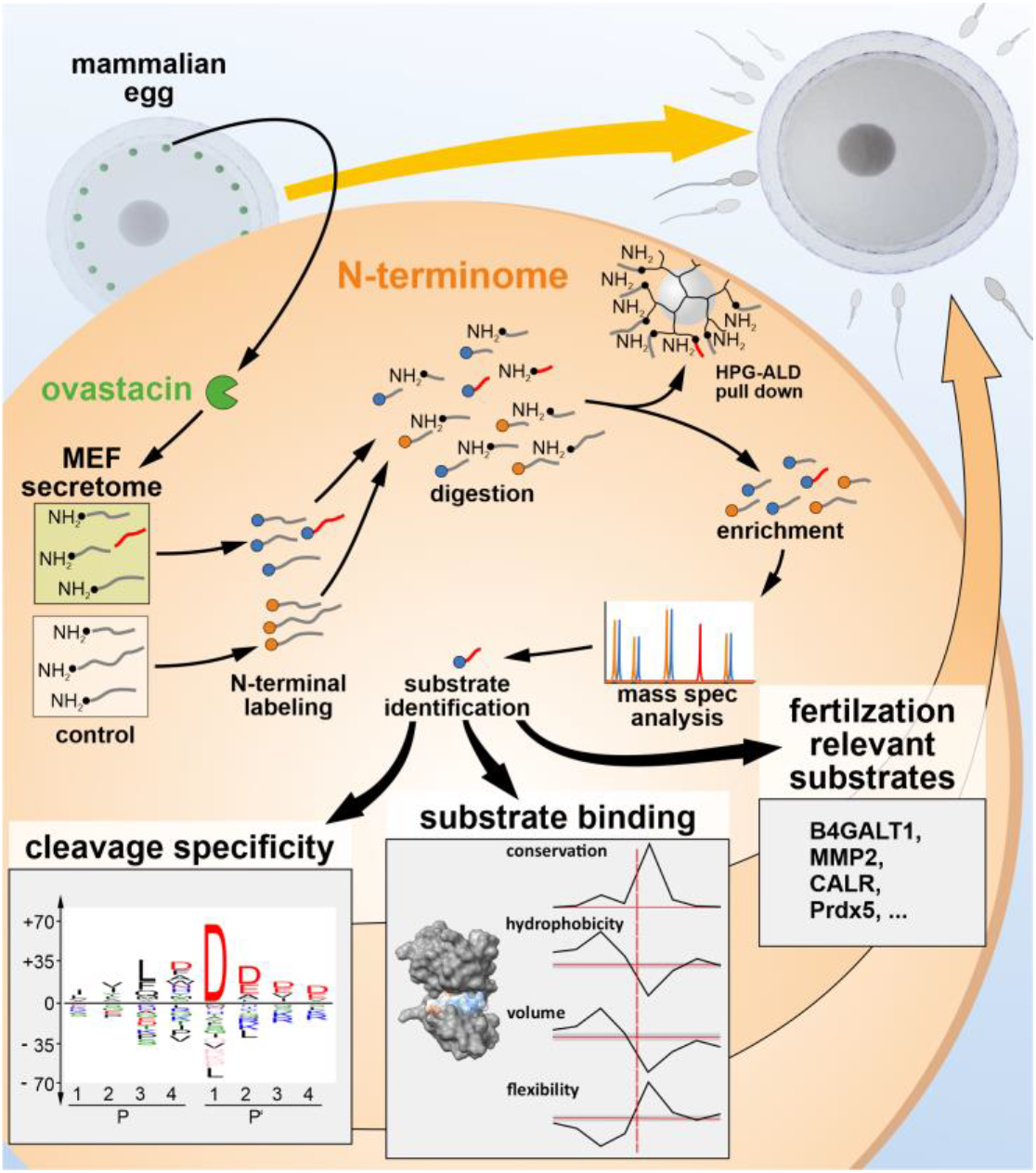

## Introduction

Several germ cell specific proteolytic enzymes have been reported to exert essential functions during egg-sperm interaction in mammalian fertilization. Among others, these include ovastacin, acrosin, testisin (TESP5, PRSS21), ADAM1b (i.e. fertilin α) and plasminogen activators (1–7), revealing a complex, coordinated and antagonist-controlled proteolytic system reminiscent of those controlling blood coagulation and fibrinolysis. However, only for two of these proteinases, namely acrosin and ovastacin, fertilization relevant cleavage sites in distinct substrates have been identified so far (1, 8–10). Considering 50 million infertile couples estimated worldwide and a rate of about 30% idiopathic infertility, 15 million couples are affected by involuntary childlessness (11, 12). Hence, there is considerable demand for further elucidation of the molecular mechanisms underlying fertilization.

Ovastacin (encoded by the gene *ASTL*), a member of the astacin family of metalloproteinases, is one of the enzymes that regulate fertilization at the level of the zona pellucida (ZP), the egg-enveloping extracellular matrix. As a typical astacin-like proteinase, ovastacin comprises a catalytic domain made up of 198 amino acid residues containing the extended zinc binding motif (H^182^ExxHxxGxxH^192^) and the strictly conserved 1,4-β-turn, called the Met-turn (M^236^), typical for metzincin proteinases (13). Additionally, it contains a unique cortical granule localization motif (D^52^KDIPAIN^59^) within the N-terminal propeptide, upstream of the conserved zinc-binding aspartate ensuring latency, and a C-terminal domain of 150 residues of unknown function (13, 14).

Upon plasmogamy and oocyte activation, ovastacin is released into the perivitelline space during the cortical reaction and cleaves a distinct peptide bond (SRLA↓ D^168^ENQ) within zona pellucida protein 2 (ZP2), thereby converting it into ZP2f (1). Thus, sperm reception is disrupted and the ZP “hardens”, which prevents sperm binding to the ZP and definitely abrogates its penetration for further sperm (1, 15). Even though ZP2 has so far been the only proven physiological substrate, cleavage of additional substrates in the complex secretome of fertilization is likely.

Phenotypically, ovastacin-deficient mice (*ASTL-/-*) are inconspicuous except for a significant reduction in fecundity (16). Here, the absence of ZP hardening results in a soft ZP, which fails to mechanically protect the embryo until implantation. During *in vitro* fertilization (IVF).

*ASTL-/-*-oocytes and embryos are covered by a vast number of bound sperm and many sperm in the perivitelline space (16, 17). However, this does not increase the rate of polyspermy. This further supports the concept that the decisive block against polyspermy is located at the oolemma level (17–19).

The absence of an endogenous ovastacin inhibitor, the plasma protein fetuin-B, leads to female infertility (20). Here small quantities of prematurely released ovastacin are sufficient to prevent fertilization. Hence, strict regulation of this proteinase is essential to maintain fertility. Conversely, supplementation of fetuin-B during IVF significantly improves the success of fertilization by effectively blocking prematurely released intracellularly activated ovastacin (16, 20, 21).

Recently, we were able to elucidate the precise mechanism of this inhibition (22–24). In mammals, only three members of the astacin family of proteinases, ovastacin and the closely related meprins (α & β) are inhibited by fetuin-B (22, 25). The underlying molecular mechanism differs greatly from the classical cystatin/papain interaction (26). This inhibition by the so called ‘raised elephant trunk’ mechanism is not due to the interaction of one or both cystatin domains of fetuin-B with the active site of ovastacin but is rather based on the warhead containing linker (C^154^PDC^157^) between the two cystatin-like domains and a hairpin loop in the second cystatin-like domain.

In this study, we utilized N-terminal amine isotopic labelling of substrates (N-TAILS) (27) on mouse embryonic fibroblast (MEF) secretome to analyze the cleavage specificity of ovastacin. We found unique characteristics of ovastacin, differing from other astacins. Furthermore, we identified a variety of potential physiological substrates including cleavage sites, which might extend the function of ovastacin during fertilization.

## Materials and Methods

### Cell culture and secretome preparation

We cultured mouse embryonic fibroblasts (MEF) in DMEM (Dulbecco’s Modified Eagle Medium (Sigma-Aldrich) with 10% FBS and 100 U/ml Penicillin / Streptomycin) (37°C, 5% CO2) in adherent culture and passaged three times per week (detaching of adherent cells with 0.05% trypsin/EDTA solution).

For preparation of conditioned cell media, MEF cells were washed three times with PBS buffer (Dulbecco’s Phosphate Buffered Saline - Sigma-Aldrich) and incubated for additional 24 h in DMEM/High Glucose (51449C, Sigma-Aldrich) with 4 mM Alanyl-glutamine (G8541, Sigma-Aldrich) and 100 U/ml Penicillin / Streptomycin). Subsequently, we collected the supernatant, treated with protease inhibitors (cOmplete™ EDTA-free Protease Inhibitor Cocktail (Merck, 04693132001) or Pefabloc® SC (CarlRoth, A154.1)), and centrifuged for 3 min at 180 x g at 4°C. The supernatant was concentrated (Vivaspin 5 kDa MWCO) to 1 – 2 mg/ml and stored on ice until further use.

### Activation of pro-ovastacin

Heterologously expressed murine ovastacin was purified and activated with human plasmin (Haematologic Technologies) as described (Karmilin *et al*., 2019; Kuske *et al*., 2021). Exemplarily, supplemental Figure 1 illustrates ovastacin purification and activation. Based on the short half-life of the ovastacin activity, the plasmin used for activation was not removed but inhibited using cOmplete™ EDTA-free Protease Inhibitor Cocktail (Merck, 04693132001) or Pefabloc® SC (CarlRoth, A154.1). The inhibition was checked via a fluorogenic assay (as exemplified in supplemental Figure 2).

### Sample preparation for Terminal Isotopic Labeling of Substrates (TAILS) analysis

The secretome of MEF cells was incubated twice with active ovastacin at a mass ratio of 100:1 for 2 h, at 37 °C or with the equal volume of activation solution without ovastacin, respectively. Subsequently, samples were reduced in 5 mM dithiothreitol (DTT), at room temperature (RT). After incubation (70 °C, 60 min) the samples were alkylated with iodoacetamide (IAA) (final concentration 15 mM, 30 min, RT, light-protected). Thereafter, proteins were precipitated with trichloroacetic acid (TCA) (28) and resuspended (200 mM HEPES, pH 7.5) to a final concentration of approx. 2 mg/ml. N-termini were labeled with light and heavy formaldehyde (^12^CH_2_O in H_2_O, and^13^CD_2_O in D_2_O (252549 and 596388, Sigma-Aldrich)), respectively. Sodium-cyanoborohydride was added to a final concentration of 40 mM, adjusted to pH 7 followed by overnight (ON) incubation at 37 °C)(29). Excess formaldehyde was captured by adjusting the sample to 50 mM Tris/HCl pH 7.0 and incubation for 4 h (37 °C). Proteins were precipitated with 9x volume acetone and 1x volume methanol (each -20 °C) ON at -80 °C and centrifuged (4500 x g for 60 min at 4 °C). The pellet was washed four times with 1 ml methanol following centrifugation (4500 x g for 15 min at 4 °C) and dissolved in 200 mM HEPES buffer (pH 8). Proteolysis into peptides was accomplished using 0.5% (w/w) Trypsin (sequencing grade; Worthington Biochemical) (37 °C, overnight). After desalting via C18 Sep-Pak columns (Waters), we enriched the labeled protein N-termini by negative selection using a polymer for enrichment (hyperbranched polyglycerol-aldehyde, HPG-ALD (University of British Columbia, Vancouver)). Peptides were adjusted to 200 mM HEPES (pH 7) and incubated with a ratio of 1:5 (w/w) of polymer and 100 mM sodium cyanoborohydride (ON at 37 °C, 40 rpm). Free aldehyde groups were saturated using 100 mM glycine pH 7 for 60 min at RT. Subsequently, the N-terminal blocked peptides were isolated using centrifugal filtration (Vivaspin 500, 10 kDa MWCO) and two washing steps (1x 300 µl 100 mM ammonium bicarbonate, 1x 300 µl 2 M guanidine hydrochloride). The samples were desalted via C18 columns and fractioned via strong cation exchange chromatography (Stagetips)(30). Afterwards, the samples were desalted again. Peptide concentrations for mass spectrometric measurement were determined by a NanoDrop ND-1000 spectrophotometer (PeqLab). Protein or peptide concentration in the course preparation was checked by Bradford- or bicinchoninic acid-assays (31, 32).

### TAILS analysis by mass spectrometry

Mass spectrometric analysis was performed using LC-MS (liquid chromatography/mass spectrometry) on a Synapt G2-S HDMS (Waters, MA, USA) in positive ion mode. Chromatographic separation of peptides was conducted on a nanoAcquity UPLC system (Waters). Peptides were separated using a C18 reversed-phase analytical separation column (HSS-T3, 75 µm x 250 mm, 1.8 µm particle size) (Waters). Samples were loaded directly onto the column (2.5 µL injection volume). Peptides were separated using a gradient of mobile phases A (water containing 0.1% (v/v) formic acid and 3% (v/v) DMSO) and B (acetonitrile containing 0.1% (v/v) formic acid and 3% (v/v) DMSO). The proportion of phase B increased from 5% (v/v) to 40% (v/v) over 90 min (300 nL/min). Afterwards, the column was re-equilibrated with phase B (90% (v/v), 600 nL/min, 5 min). Total measurement time was 120 min. Data were acquired using HD-DDA (33). [Glu^1^]-fibrinopeptide was used as lock mass at 100 fmol/µL and sampled every 30 s into the mass spectrometer via the reference sprayer of the NanoLockSpray source. Processing of raw data and database searching was performed using PEAKS (version 8.5; https://www.bioinfor.com/) (34–36). We reviewed liquid chromatography plots individually to exclude false positive results. Single signals (FDR = 0.01) identified by PEAKS were checked for the presence of a double signal caused by 6 Da mass difference due to dimethylation, which was considered a duplicate and excluded from the data set. Peptides with naturally acetylated N-termini as well as peptides with unlabeled N-termini or double labeled with light and heavy formaldehyde were also excluded. All peptides identified by PEAKS that had N-terminal labeling by heavy formaldehyde and no analogue with N-terminal labeling by light formaldehyde were manually selected. Samples were prepared as two independent biological replicates and each analyzed in technical triplicates. The final data set only included peptides that could be assigned to a single UNIPROT accession number from *Mus musculus*. Amino acid residues N-terminally of the cleaved peptide bond (i.e. “nonprime side”) were determined by bioinformatic alignment of the sequences with the UNIPROT database. Cleavage sites less than 10 residues distant from N- or C-terminus were considered.

Comparing all identified cleavage sites with aspartate in position P1’ (cleavage preference of astacin proteases against the full data set revealed a negative correlation with lysine and arginine in position P1 (tryptic cleavage specificity). Therefore, in order to exclude peptides generated e.g. by not entirely inactivated plasmin carried over from ovastacin activation, the data sets from both biological replicates were further subjected to cluster analysis (GibbsCluster 2.0) (37, 38). This is exemplified for one of the replicas in supplementary figures 3 and 4. The identified clusters from both biological replicates, with the typical acidic residues on the prime side (39), were combined for the further analyses. The dataset was limited to proteins within the physiological context of the extracellular space by analysis using the DAVID Bioinformatics database (40, 41).

### Identification of the cleavage specificity and physicochemical properties of ovastacin

Determination of cleavage preferences derived from the substrate cleavage sites, their representation as sequence logos, and statistical variations in the abundances of residues was performed using iceLogo (42). In order to equally compare cleavage preferences of other proteases based on the cleavage sites annotated in the MEROPS database (10), only cleavage sites with all positions from P4 to P4’ identified were used. The “subLogo” function was used to check for cooperativity between residues in different positions. To calculate the degree of conservation of residues in each position the “Conservation Line” function with a BLOSUM62-substitution matrix (43) was used.

For determination of parameters needed for evaluation of physicochemical properties the “Aa Parameter” function was used. Underlying parameters are listed in the AAindex1 database (http://www.genome.jp/aaindex/). The “Transfer Energy, organic solvent/water” template was used to determine hydrophobicity (44), the “Residue Volume” template to determine residue volume (45), and the “Average flexibility indices” template to determine residue flexibility (46).

### *In silico* structure prediction

Models were visualized using Chimera 1.13.1 software. Homology models were calculated using Modeller (https://salilab.org/modeller/) (47, 48) via the Chimera interface. Alignments necessary for generating homology models, were also performed with Chimera (via BLOSUM-62 matrix (43), gap opening penalty 12 / gap extension penalty 1). For model creation with more than one template, the structure-specific secondary structure score was employed. The homology model with most favorable (negative) zDOPE (Discrete Optimized Protein Energy) value was selected.

The stereochemical quality of the models generated was evaluated using the PROCHECK software (www.servicesn.mbi.ucla.edu/PROCHECK) (49, 50). Interactions between ovastacin and identified substrates were simulated *via* the ClusPro software (https://cluspro.org) (51–53) using the generated models. From all calculated values, those 1000 with the most favorable (lowest) energy were selected and searched for clusters in the configurations.

## Results

### TAILS Analysis

We performed two N-TAILS analyses on native MEF secretome with heterologously expressed murine ovastacin to identify potential physiological relevant substrates of ovastacin, as well as its cleavage preference. Together, a total number of 855 unique cleavage sites were identified in 489 proteins. After restriction to the physiological context, the extracellular space, by database mining, we identified a dataset of 353 unique cleavage sites in 163 proteins (Supplemental Table S1 and Supplemental Table S2).

### Ovastacin features the typical acidic prime side specificity of the astacin metalloproteinase family

Results are presented according to the concept by Schechter and Berger (54). Peptide residues N-terminal to the cleavage site are indicated as nonprime (P) and C-terminal as prime (P’). The summary of the cleavage profile of the 855 unique cleavage sites exhibits very high similarity to the only physiologically verified cleavage site SRLA↓ D^168^ENQ in ZP2 (1)(Figure 1). Ovastacin significantly prefers negatively charged residues in positions P1 to P6’, especially for aspartate in positions P1’ (+70.1%, rel. to natural occurrence) and P2’ (+16.6%). No strong preference is evident for position P1. Nevertheless, aspartate (+7.6%), phenylalanine (+4.8%), alanine (+4.6%), tryptophan (+4.0%), asparagine (+3.7%) and histidine (+3.5%) are found predominantly. In position P2, leucine (+19.8%), phenylalanine (+8.2%) and glutamine (+5.4%) exhibit the highest abundance. Hydrophobic residues are preferred in the range from P4 to P1 with aromatic residues at higher frequency in positions from P3 to P1. On the prime side, on the other hand, acidic residues, especially aspartate, predominate in the range from P1 to P6’. This is highlighted by the graphic iceLogo representation (figure 1 D). Ovastacin cleavage sites in substrates reveal a strong cleavage preference for negatively charged residues on the prime side, especially for aspartate in position P1’. In general, aspartate is clearly favored over glutamate in all positions of the prime side. On the nonprime side, leucine, phenylalanine and methionine are particularly preferred in position P2, and tyrosine, valine, isoleucine and phenylalanine in position P3.

**Figure 1.**
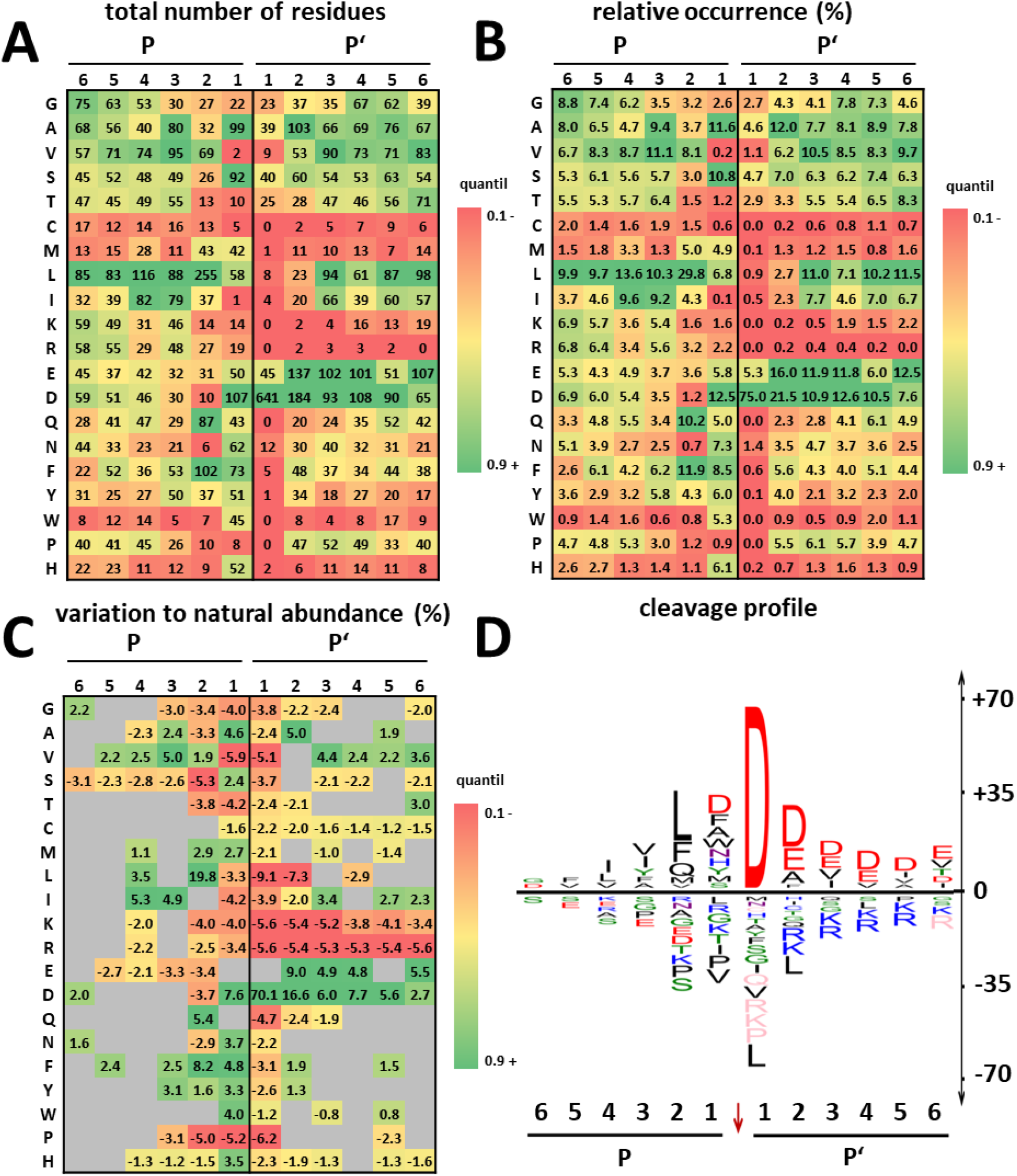
Heatmap and iceLogo of cleavage sites (n= 855) identified via N-TAILS. Residues are normalized to their natural occurrence in the mouse proteome (*Mus musculus*). Distribution of residues in positions P6-P1 (nonprime side) and P1’-P6’ (prime side). Displayed for each position are the total number of residues (A), their relative abundance in percent (B) and the difference to the natural abundance in percentage points as heatmap (C) and as iceLogo (D). The deviations were determined with iceLogo (42). Positions without significant changes (p>0.05) are depicted in gray for the heatmaps, the iceLogo displays only significant changes (p≤0.05). Amino acids abbreviated as single letter.

### Physicochemical properties of ovastacin compared to other astacin proteinases

The cleavage preference determined for ovastacin (figure 1) shows high agreement with the cleavage characteristics of other astacins (BMP1, meprin α, meprin β, and LAST MAM) partly due to the preference for the negatively charged aspartate residue in position P1’ (39). To identify ovastacin-specific characteristics, we analyzed in detail the physicochemical properties of cleavage sites, i.e electrostatics, hydrophilicity/hydrophobicity, volume, flexibility and degree of conservation of distinct residues flanking the cleavage site. All signatures of the individual astacin members (supplementary figure 3) feature the highest degree of conservation in position P1’ of the cleavage site, and to lesser extent in the positions P2 to P2’ for meprin α only. The hydrophobicity of the residues reaches its minimum in positions P1 and P1’, respectively, and its maximum in positions P3 and P2. The side chains with the largest volumes tend to be in positions P3 to P2, those with the smallest in the range P1 to P2’. The flexibility of the residues is lowest between P3 to P2 and its highest in the positions P1’ to P2. Therefore, the typical substrate of the astacins may be characterized as follows: non-hydrophobic, small and flexible residues in position P1’ and to a slightly lesser extent in the flanking positions P1 and P2’, and larger, non-flexible hydrophobic residues in positions P3 to P2. All investigated astacins exhibited a common pattern, despite significant differences in the individual substrate sequences, in the parameters studied. Therefore, these are combined in a single signature of astacins (figure 2 A). Compared directly with the other astacins, ovastacin (figure 2 B, C) exhibits a particularly high degree of conservation of residues in position P1’ and, to a lesser extent, in position P2. The hydrophobicity in positions P4 to P1 (especially in P2) is even more pronounced, while position P1’ is rather hydrophilic. Furthermore, the volume of side chains on the nonprime side, especially in position P2, is larger than for the other astacins. The flexibility of the residues in ovastacin substrates is comparatively lower in the region P4 to P1, but significantly higher in position P1’. Among astacins, the physicochemical cleavage signature of ovastacin shows the greatest similarity to that of meprin β (supplementary figure 3 C). However, in direct comparison, meprin β significantly favors glutamate in position P1’.

**Figure 2.**
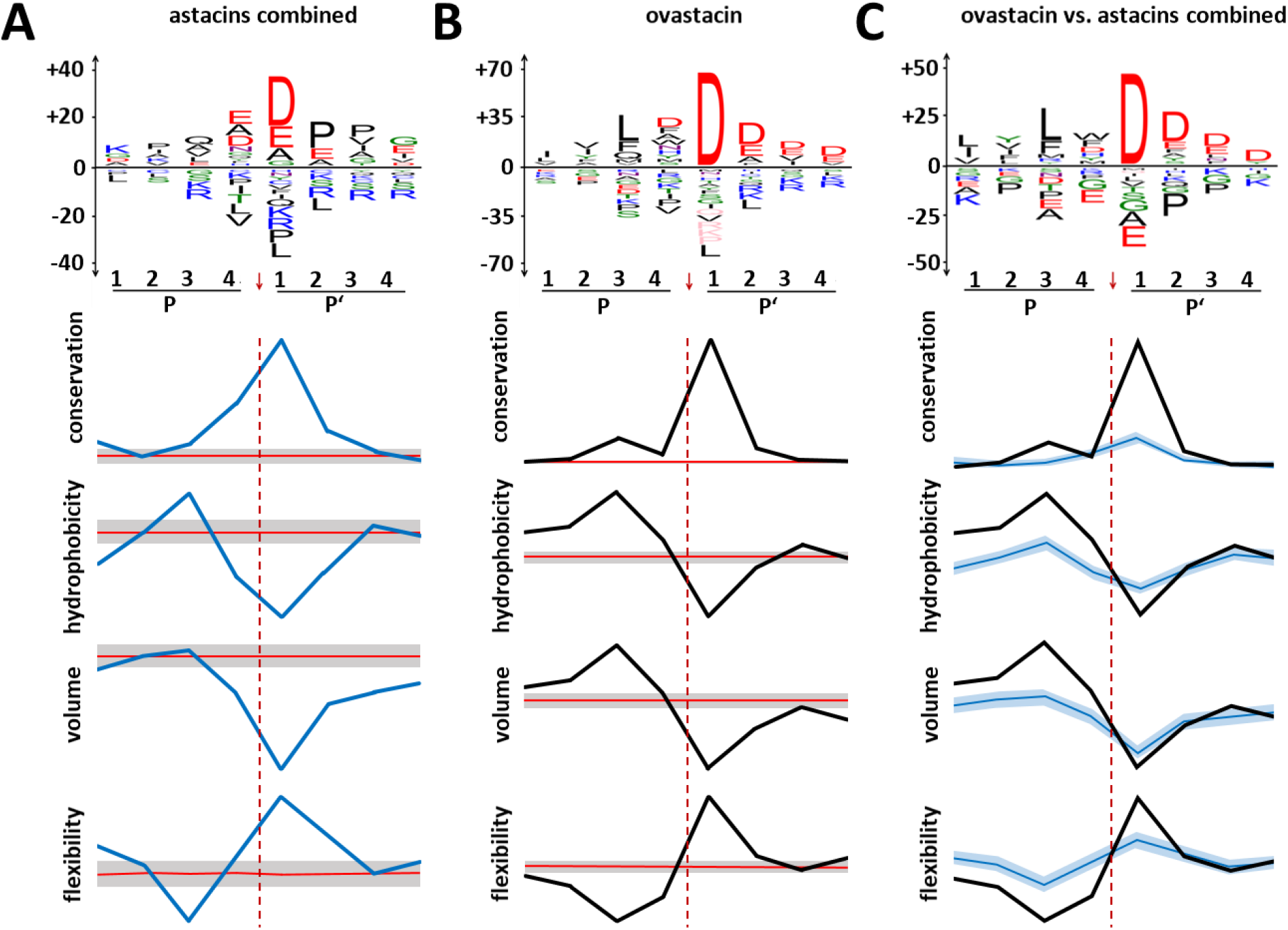
Physicochemical properties of astacin proteases and ovastacin. Overview of the common cleavage preferences and physicochemical properties of all studied astacin proteases combined (left), ovastacin only (middle) and comparatively of all astacin proteases versus ovastacin (right). The analysis was performed on the basis of the cleavage sites deposited in the MEROPS database (astacin (P07584), n=199; BMP1 (P13497), n=25; LAST_MAM (B4F320), n=415; LAST (B4F319), n=76; meprin α (Q16819), n=700; meprin β (Q16820), n=879) and ovastacin (Q6HA09), n=855 (this study). Depicted in each case is a sequence logo (upper panel) displaying the difference to the natural abundance of residues in the cleavage site positions P4-P4’, the degree of conservation of positions according to the BLOSUM62 substitution matrix (43) (upper graph), the hydrophobicity of the residues (44) (second graph from the top), the volume of the residues (45) (second graph from the bottom) and the flexibility of the residues (46) (bottom graph). The significance level is 95%. The error range is displayed in gray/light blue. The diagrams allow a semi-quantitative evaluation, since iceLogo does not allow scale normalization.

### Substrate interaction

We first created homology models (supplementary figure 4) to analyze the substrate interaction with the catalytic domain of ovastacin based on the physicochemical properties of the surface. The model with the combined templates of ZHE-1 (PDB 3LQB(55)) and meprin β (PDBs 4GWN and 4GWM (56)) was used for this purpose. Based on the observation in standard orientation (57), positive charges are primarily located on the upper rim of the right half of the catalytic cleft (figure 3 A1). The bottom of the catalytic cleft displays non-charged hydrophilic properties, except for a hydrophobic region in the left periphery (figure 3 A2). Hydrophobic regions stretch across the upper and lower rim of the active side cleft. Strongly hydrophobic regions are located at the left margin of the catalytic cleft as well as at its basis. In summary, there is a transition in the physicochemical properties from strongly hydrophobic to strongly hydrophilic with positive charges in the active side cleft, in standard orientation from left to right (i.e. from nonprime to prime side). Arginine, lysine and histidine are responsible for the formation of the hydrophilic, strongly positively charged regions in the S’ region (figure 3 B). Also displayed are the large, rigid, and hydrophobic residues leucine, isoleucine, valine, phenylalanine, tryptophan, methionine and tyrosine (figure 3 D). Evaluation of the cleavage sites obtained from N-TAILS analyses revealed V-L-D↓ D-D-D as the theoretically most preferred sequence spanning positions P3 to P3’ in ovastacin cleavage sites (figure1 D). This peptide was docked virtually into the catalytic cleft of ovastacin using Chimera (figure 3 C). It is apparent that the aliphatic substrate residues Val (P3) and Leu (P2) are in the region between Ile216, Ile219 and Phe216 on one side, Phe54 on the other side and Trp191 at the base of the active side cleft (figure 3 C). The aspartate residues of the prime side are, depending on the exact orientation of the side chains, proximate for interacting with the sidechain of Arg264. Arg174 appears too distant for a direct interaction at least for P1’ and P2’. Likewise, Lys175 is too distant for a direct interaction with the aspartate residues in position P1’-P3’.

**Figure 3.**
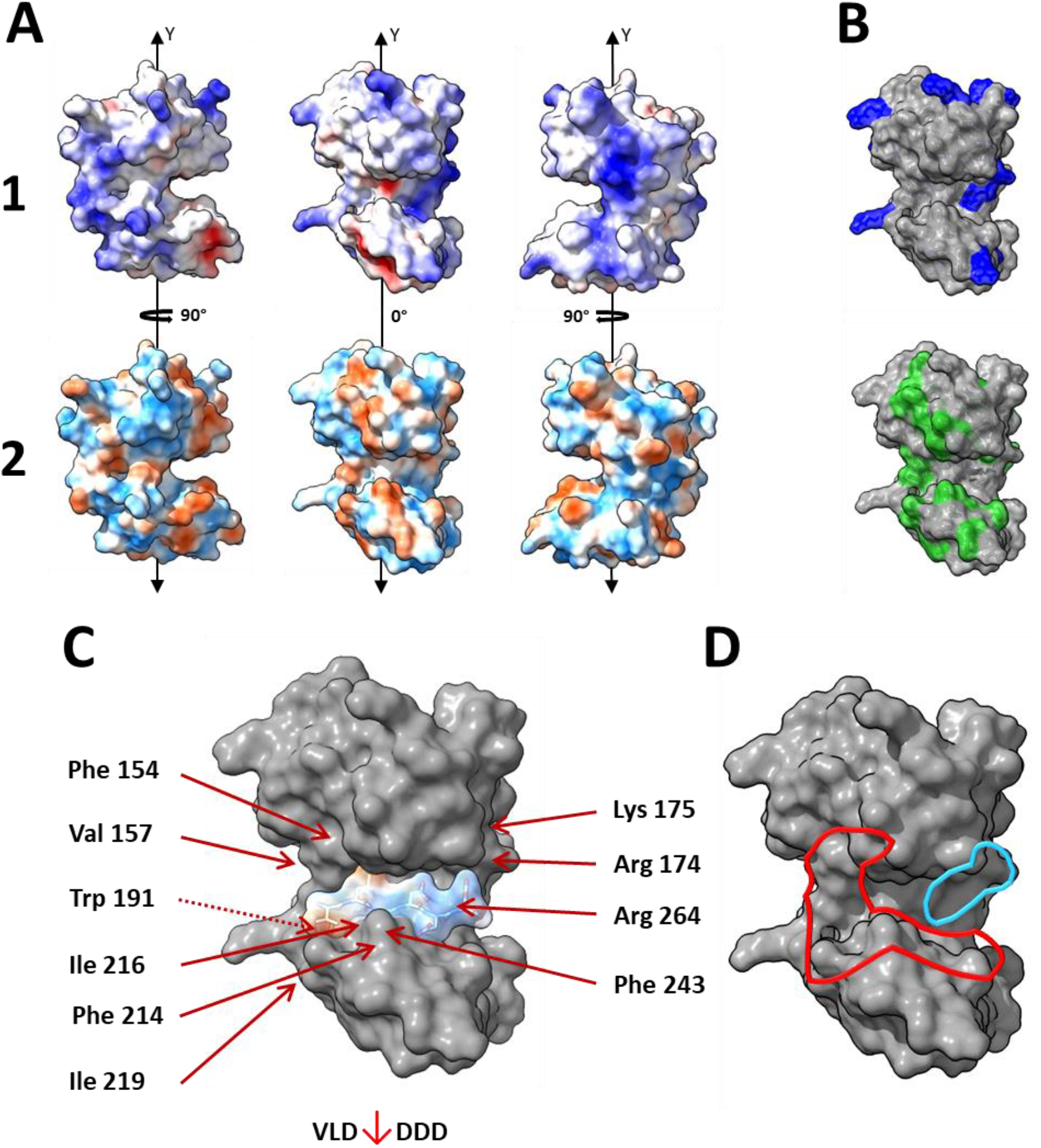
Structural analysis of substrate interaction. Surface models of the catalytic domain of murine ovastacin (Q6HA09; position 92-282) created with Modeller (47) using the templates of the catalytic domains of meprin β and ZHE-1 (PDBs: 4GWN, 4GWM, and 3LQB). (A 1) Coulomb potential of the surface highlighting areas with positive (blue, 10 kcal/(mol*e)) and negative (red, -10 kcal/(mol*e)) electrostatic potential. (A 2) Surface areas of hydrophilic (light blue) and hydrophobic (orange) residues according to the Kyte-Doolittle scale (74). (B) Upper model with all accessible positively charged residues (Arg, Lys) and lower model with all accessible hydrophobic residues (Leu, Ile, Val, Pro, Phe, Trp, Met and Tyr). (C) Synthetic peptide (surface areas of hydrophilic (light blue) and hydrophobic (orange) residues) according to the most preferred residues in position P3 to P3’ (figure 1) and inserted into the catalytic cleft via Chimera; highlighted residues of ovastacin are discussed (D) Hydrophobic (outlined in red) and hydrophilic (outlined in blue) regions in the catalytic cleft.

### Physiological relevant substrates identified via TAILS analysis

Cleavage of ZP2, the only confirmed physiological substrate of ovastacin so far, is a key regulatory event in fertilization since it destroys the sperm receptor and mediates ZP-hardening. By screening the 489 substrates cleaved by ovastacin in our N-TAILS analysis for proteins involved in fertilization, we identified 12 different potentially fertilization-relevant proteins including 29 cleavage sites (Table 1 & 2). Several examples are to be mentioned explicitly. The β-1,4-galactosyltransferase (B4GALT1) is localized on the sperm membrane and binds sugar side chains of ZP3, thereby mediating additional attachment to the ZP (58). Calreticulin (CALR3) is localized on the surface of the acrosomal region of sperm (59) and also on the oolemma (60). There is evidence for its involvement in oocyte signal transduction (61) and the establishment of the polyspermy block after its exocytosis from the oocyte (62). Chaperonin containing TCP1 (Cct2) localizes also on the sperm surface and is directly involved in the binding of sperm to oocytes (63, 64). Matrix metalloprotease 2 (MMP2) is associated with the acrosome and potentially contributes to the penetration of the ZP (65). Taken together, detection of cleavages in these proteins strongly indicates a more complex function of ovastacin within the proteolytic web of egg-sperm interaction beyond regulation of ZP hardening.

**Table 1.** Search terms for identification of substrates associated with fertilization. Search terms in the DAVID bioinformatics database (40, 41) including their category, the p-value, the number of substrates in the dataset and their fraction of the total dataset.

| search term | category | P-value | substrates n | proportion [%] |
| --- | --- | --- | --- | --- |
| *zona pellucida* receptor complex | GOTERM_CC_DIRECT | 3.5 E-3 | 4 | 0.8 |
| binding of sperm to *zona pellucida* | GOTERM_BP_DIRECT | 1.4 E-2 | 5 | 1.0 |
| estrogen signaling pathway | KEGG_PATHWAY | 7.4 E-2 | 8 | 1.6 |
| *in vitro* fertilized eggs | UP_TISSUE | 1.4 E-2 | 13 | 2.7 |

## Discussion

Our N-TAILS analysis identified hundreds of ovastacin substrates and revealed the substrate specificity of ovastacin. In fact, ovastacin shares a high degree of similarity with the substrate specificity of meprin β. Accordingly, some well-known meprin β substrates, such as the amyloid precursor protein (APP), were cleaved by ovastacin as well (Supplemental Table 1). However, due to the spatio-temporal restriction of ovastacin to oocytes or the functional sphere of the egg-sperm interaction, these are presumably of no physiological relevance. The restriction to substrates to the extracellular space or the context of fertilization reduced the number of physiologically relevant ovastacin substrates in the present study from 489 to 163 and twelve, respectively. Owing that ovastacin expression is extremely limited to oocytes, this rigorous restriction may be appropriate (66). Although our data suggest a physiological processing, further steps are required to provide the physiological evidence. At least, a confirmation of the data obtained in a cellular context, e.g. egg or sperm secretome, is required to verify the physiological aspects of a cleavage by ovastacin. The germ cell specific expression pattern might compromise the activity or substrate selection of ovastacin e.g. by potential pre-processing or interactions of substrates. This would also explain why we were not able to identify the only physiologically proven substrate. ZP2 is not expressed by MEF cells (67). Establishment of the N-TAILS approach on germ cell secretome would remedy these problems.

Most of the fertilization-relevant substrates listed in Table 2 have been found to positively influence sperm viability or sperm-oocyte binding (63–65, 68, 69). If cleavage of these substrates results in their loss of function, ovastacin could be the initiator of a multifactorial regulation of sperm interaction. For instance, β-1,4-galactosyltransferase (B4GALT1) mediates the binding of sperm head to oligosaccharides of ZP3 until exocytosis of the acrosome (70). Although this interaction is not essential, unlike sperm binding to ZP2, its function is poorly understood. Proteolysis of B4GALT1 at S^248^ or D^249^, respectively, in a loop region between two central β-sheets, is likely to alter the conformation and binding properties to ZP3.

**Table 2.**
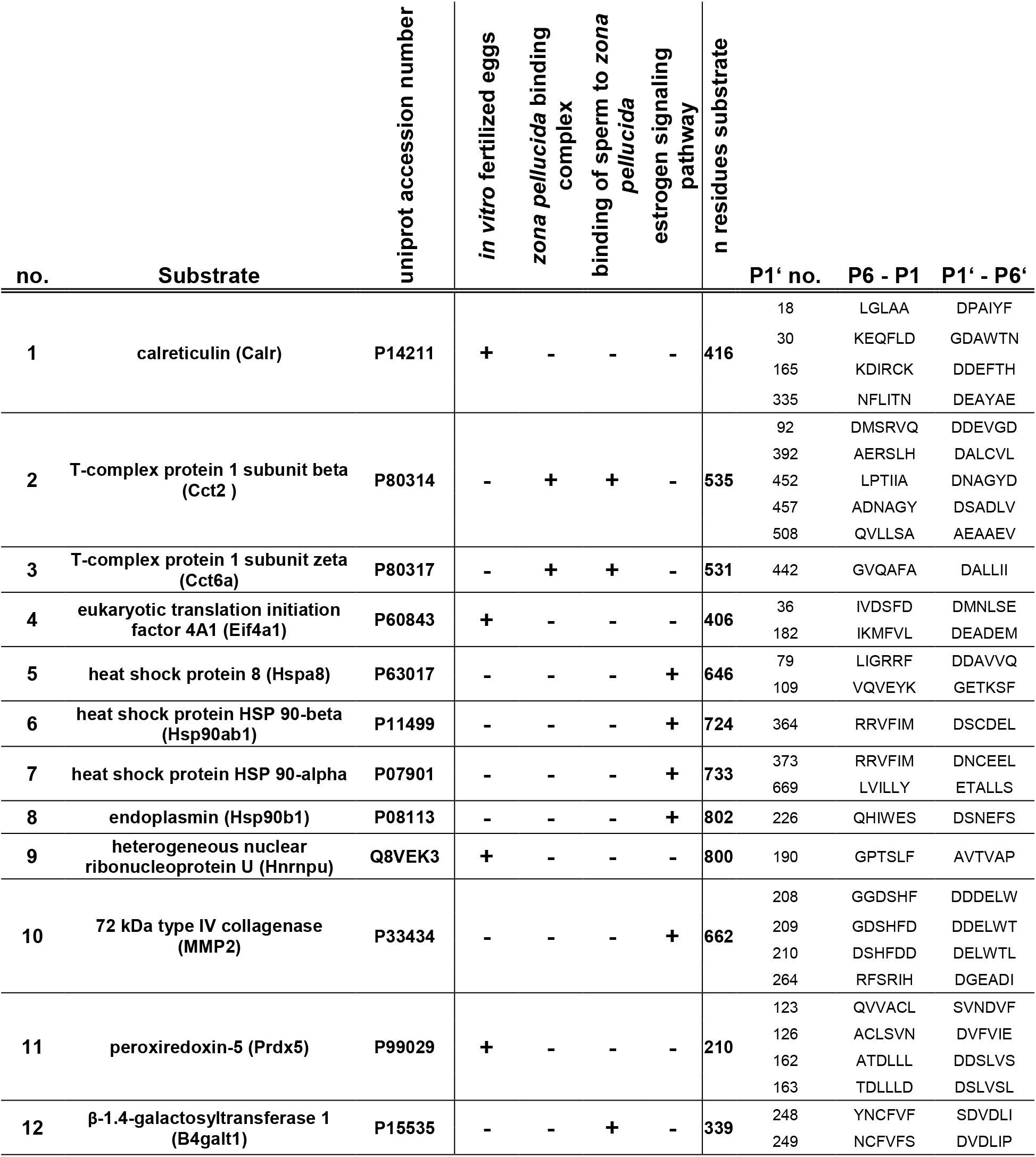
Substrates of ovastacin with proposed function in fertilization. In TAILS analysis identified substrates whose annotations in the DAVID Bioinformatics database suggest a potential physiological impact on the fertilization. Listed are the substrates, the UniProt accession number, the search term, the total number of amino acids, the cleavage site position and the corresponding residues on the nonprime and prime side.

Similarly, for instance, the D^208^, D^209^ and D^210^ in MMP2 are in the β-sheet above the zinc-binding active site helix (Table 2). From there, the chain enters the first fibronectin-like domain, which is an important exosite for binding protein substrates. Hence, cleavage within this region would most likely disrupt (protein-) substrate binding to MMP2 and abrogate its function and thereby may impair the penetration capacity of sperm. Reduction of sperm viability and hindrance of sperm binding to the oocyte would extend the effect of the already known ZP hardening by ovastacin. Recently, the regulation of ovastacin activity has been shown to be clinically relevant in humans (21, 71, 72). Thus, these potential substrates are of major interest for this field.

The cleavage preferences identified in this study displayed a clear preference for aspartate in position P1’ and, to a lesser extent, for aspartate and glutamate in the P2’ to P4’ region of the prime side. In addition, there is a preference for leucine in position P2 of the nonprime side. A similar preference for aspartate in position P1’ has been also described for the astacin proteases BMP1, meprin β, and LAST MAM (MAM-domain containing astacin from the horseshoe crab *Limulus polyphemus*) (supplemental figure 3). The substrate binding was analyzed by modeling. This interaction might be mediated by interactions with Arg174 and Arg264 and by the properties of hydrophobic (aromatic as well as aliphatic) residues of the nonprime side. In standard orientation, Phe154 is located at the upper left edge of the catalytic cleft, Val157 and Trp191 are in the bottom section, and Ile216, Ile219, Phe214 and Phe243 are present at the lower left edge. Together, these residues wrap around the peptide chain of the substrate in the left half of the catalytic cleft in a cuff-like fashion.

Crayfish astacin and LAST, unlike ovastacin, do not display preference for negatively charged residues in position P1’. However, we found great similarities between them in terms of distribution of physicochemical properties across the different positions relative to the cleavage site. All analyzed astacins share similar physicochemical properties in their substrate binding clefts. Thus, based on these findings, the different astacin proteases can be designated as variants of a common prototype. Accordingly, this prototypical signature is independent of the substrate sequence preferred by the respective enzyme. Due to the limited data currently available, this analysis was performed for only seven astacins (astacin, BMP1, LAST, LAST MAM, meprin α, meprin β, and ovastacin (Supplemental figure 3)). However, this analysis of physicochemical properties allows for deeper understanding of cleavage preference and substrate selectivity of proteases in general. This could contribute to higher precision when predicting potential protease substrates, as well as to improvement in the design of proteases with a specific substrate specificity.

The knowledge of the biochemical properties that distinguish the cleavage of substrates by ovastacin from the closely related meprins could be useful for the development of more specific and selective inhibitors. Such inhibitors would significantly reduce the amount of oocytes required for IVF (21, 73). Additionally, these could overcome the problems of proteinogenic inhibitors (e.g. half-life, application restrictions, production) such as fetuin-B.

## Conclusion

In summary, using N-TAILS, we were able to identify new substrates of the metalloproteinase ovastacin with potential physiological relevance. This deepens insight into the complexity of mammalian fertilization. The cleavage preference of ovastacin towards these native substrate proteins was resolved and resulted in the recognition of a globally applicable cleavage site signature based on physicochemical properties. This signature might serve as a useful tool to elucidate protease-substrate interactions and provide broader explanations for the genesis of cleavage preference in proteases.

## Data availabilitystatement

The LC-MS/MS data have been deposited to the ProteomeXchange Consortium via the JPOST repository with the dataset identifier PXD038561 (ProteomeXchange) and JPST001944 (jPOST).

## Acknowledgments

The MEF cell line was kindly provided by Oliver Schilling (University Medical Center, Freiburg, Germany). We thank Michael Plenikowski for preparation of the graphical abstract.

## Author Contributions

Substantial contributions to conception and design (MF, UD, WS, HK). Acquisition of data, or analysis and interpretation of data (MF, UD, MŁ, NvW, WS, HK). Drafting the article or revising it critically for important intellectual content (MF, NvW, ST, CB, WS, HK). Final approval of the version to be published (all authors).

