## Supplemental Data for "Substrate profiling of the metalloproteinase ovastacin – Implications for its physiological function in mammalian fertilization"

Running title: Ovastacin – Substrate profiling

**This PDF file includes:** Supplemental Figures 1 to 6

**Supplemental Figures and Tables**

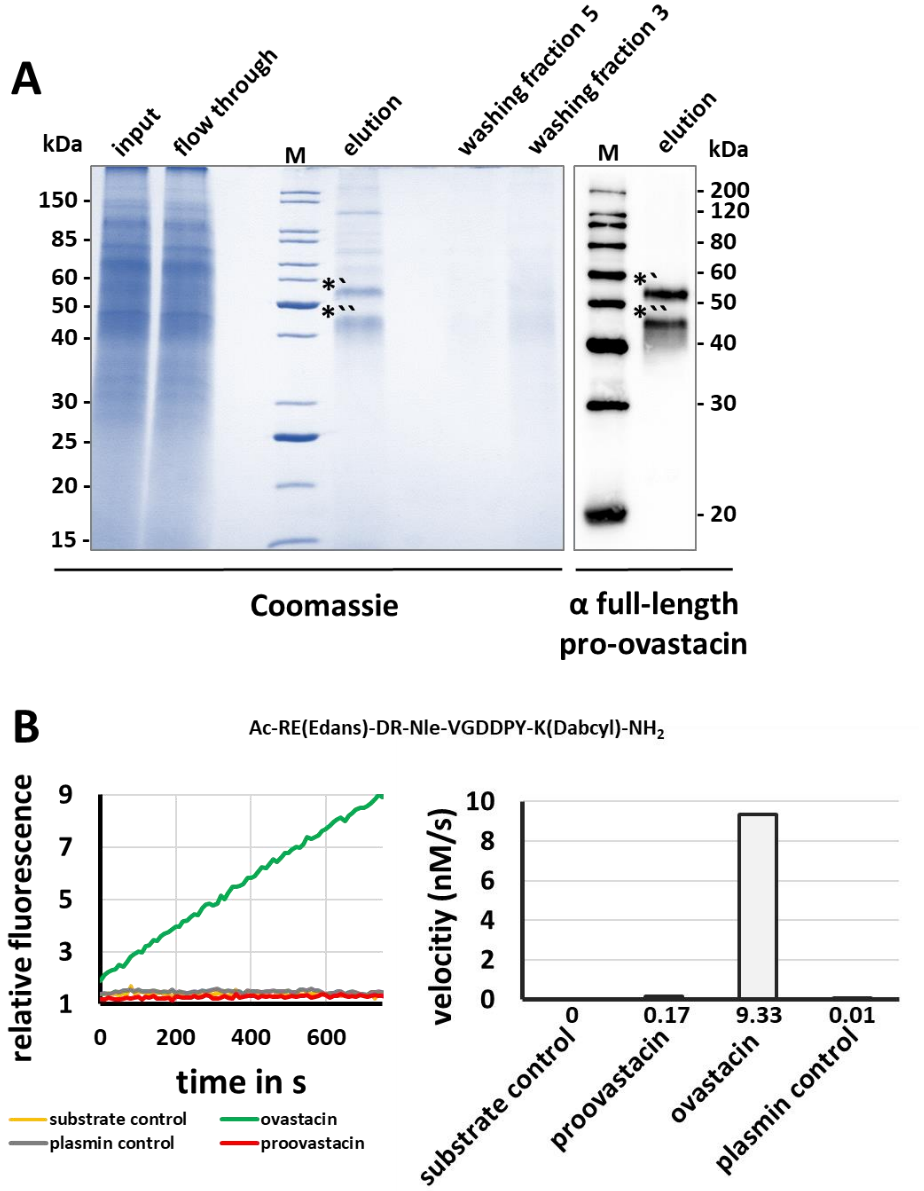


**Supplemental Figure 1. Purification and activation of ovastacin.** (A) Separation of the samples by reducing 12% SDS-PAGE stained with Coomassie brilliant blue (left) or immunoblot with an anti-proovastacin antibody (right). From left to right: input (secretome of ovastacin expressing High Five cells), flow through (of Strep-Tactin affinity chromatography), protein marker, Strep-Tactin elution fraction (proovastacin) and washing fractions (of Strep-Tactin affinity chromatography). *`full-length pro-ovastacin, *`` C-terminally truncated pro-ovastacin. (B) Fluorogenic activity assay with 25 µM Ac-RE(Edans)-DR-Nle-VGDDPY-K(Dabcyl)-NH2; using 200 nM proovastacin (0.17 nM/s), plasmin (20nM) activated 200nM ovastacin (9.33 nM/s), plasmin control (0.01 nM/s) in 50mM Tris, 150mM NaCl 0,01% Brij-35, 2 mg/ml cOmplete™ EDTA-free Protease Inhibitor Cocktail, pH7,4; Incline of relative fluorescence over time (left), velocity (right).

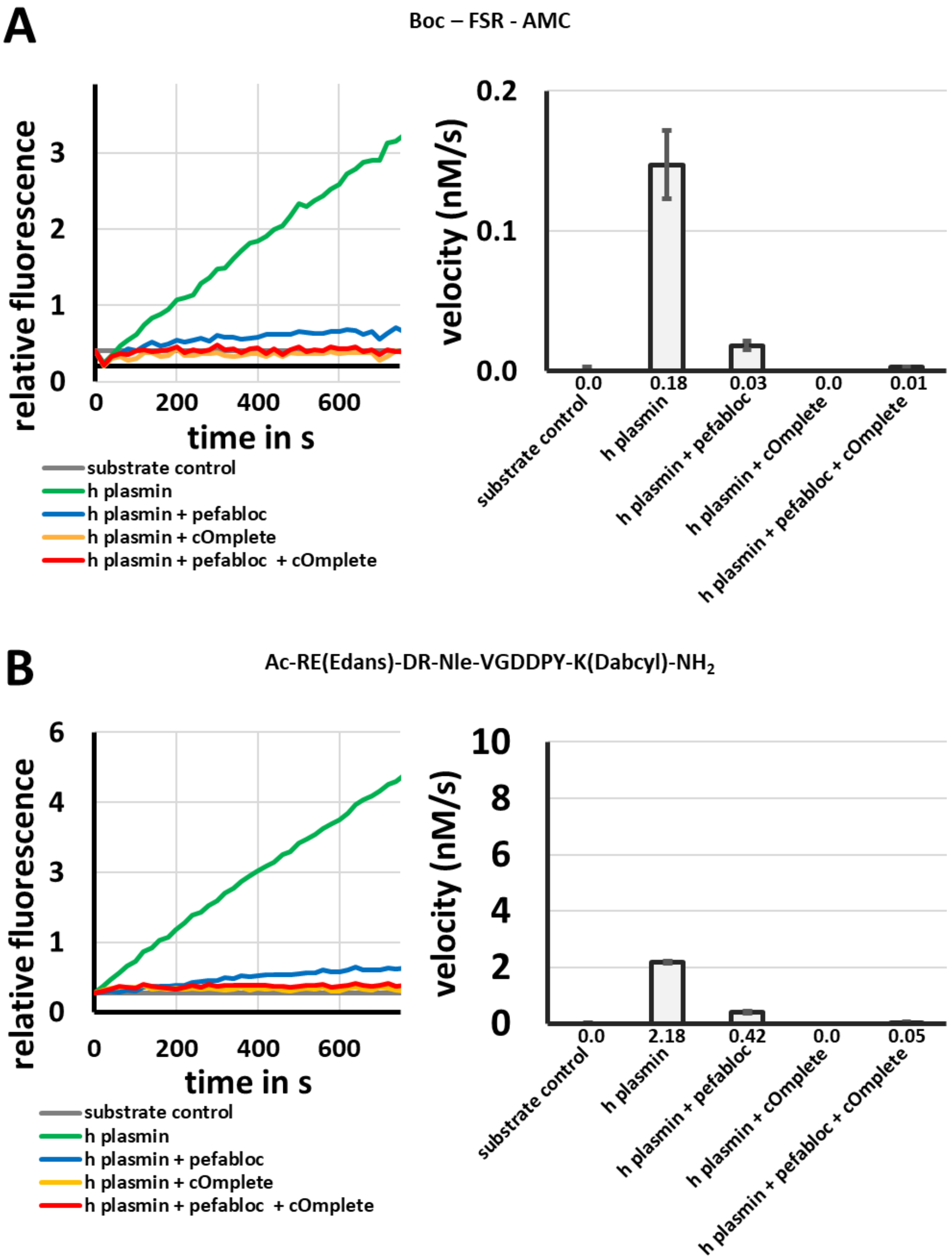


**Supplemental Figure 2. Evaluation of plasmin inhibition.** (A) Fluorogenic activity assay with 20 µM Boc-FSR-AMC using 5 nM plasmin treated as indicated with Pefabloc® SC (10 mM) and/or cOmplete™ EDTA-free Protease Inhibitor Cocktail (2.5 mg/ml) (B) Fluorogenic activity assay with 25 µM Ac-RE(Edans)-DR-Nle-VGDDPY-K(Dabcyl)-NH2 using 5 nM plasmin treated as indicated with Pefabloc® SC (10 mM) and/or cOmplete™ EDTA-free Protease Inhibitor Cocktail (2.5 mg/ml). Incline of relative fluorescence over time (left), velocity (right).

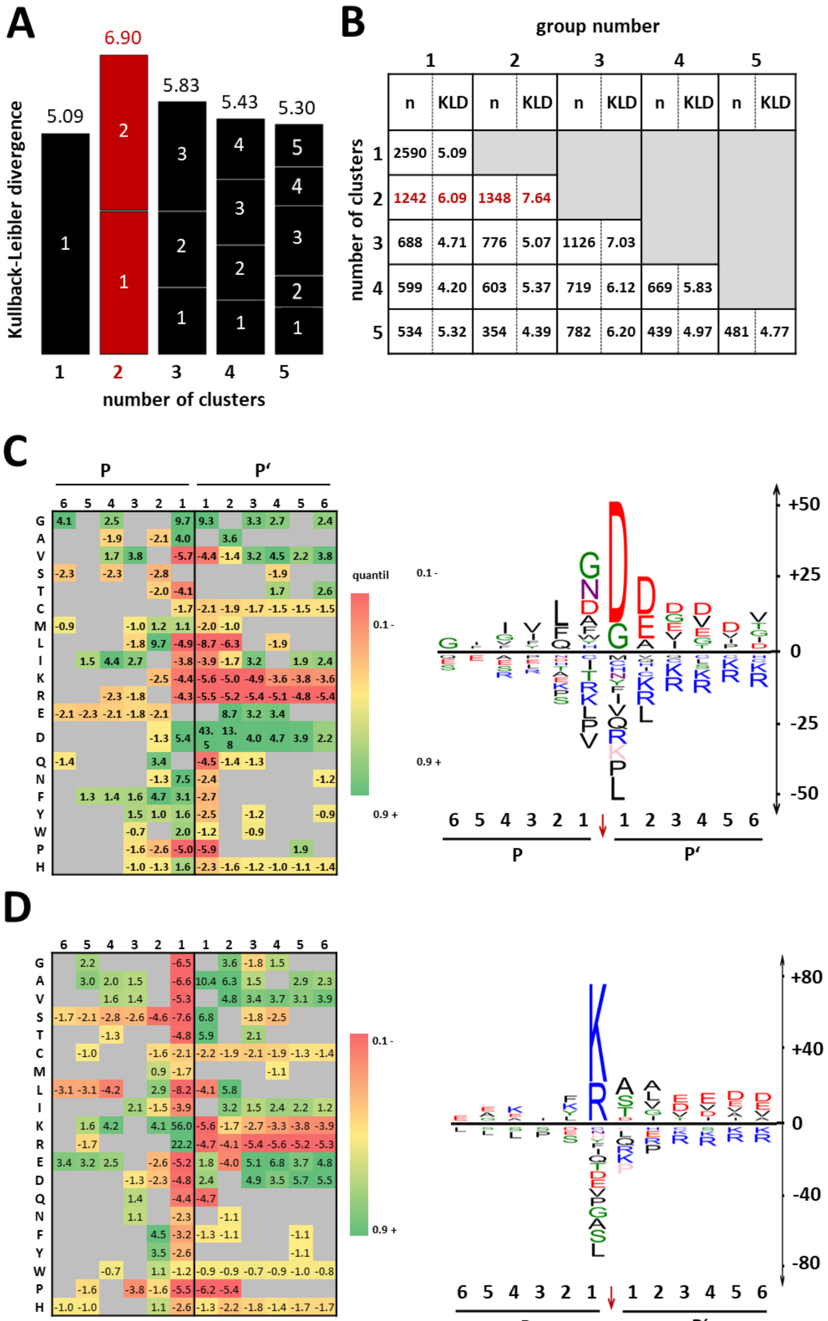


**Supplemental Figure 3. Cluster analysis of cleavage sites identified via N-TAILS Part 1.** Cluster analysis using the GibbsCluster-2.0 server (37, 38) based on the alignment of all cleavage sites identified (A) Displays the average Kullback-Leibler divergence (KLD) for the calculations with 1 to 5 clusters. (B) Shows the number of cleavage sites (n) for each cluster, as well as their Kullback-Leibler divergence (KLD). Identified subgroups 1 (C) and 2 (D) are displayed as iceLogo and heatmap, respectively (42). Shown are significant deviations ($p \leq 0.05$) from natural occurrence in percentage points. Parameters used for cluster analysis: motif length: 12, number of clusters 1-5, $\lambda=0.8$. Amino acids abbreviated as single letter.

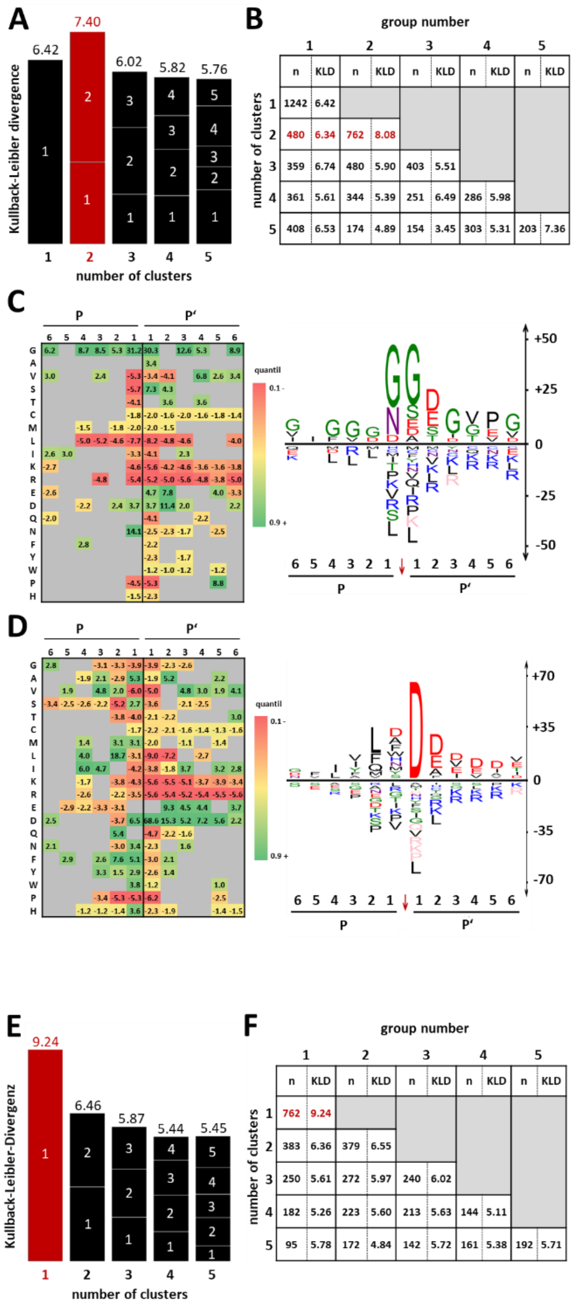


**Supplemental Figure 4. Cluster analysis of cleavage sites identified via N-TAILS Part 2.** Cluster analysis of subgroup 1 (supplemental figure 3) using the GibbsCluster-2.0 server (37, 38) based on the alignment of all cleavage sites identified (A) Displays the average Kullback-Leibler divergence (KLD) for the calculations with 1 to 5 clusters. (B) Shows the number of cleavage sites (n) for each cluster, as well as their Kullback-Leibler divergence (KLD). Identified subgroups 1 (C) and 2 (D) are displayed as iceLogo and heatmap, respectively (42). (E) and (F) display the results of cluster analysis of subgroup 1 (see (A)). Shown are significant deviations ($p \leq 0.05$) from natural occurrence in percentage points. Parameters used for cluster analysis: motif length: 12, number of clusters 1-5, $\lambda=0.8$. The amino acid single letter code is used for abbreviation.

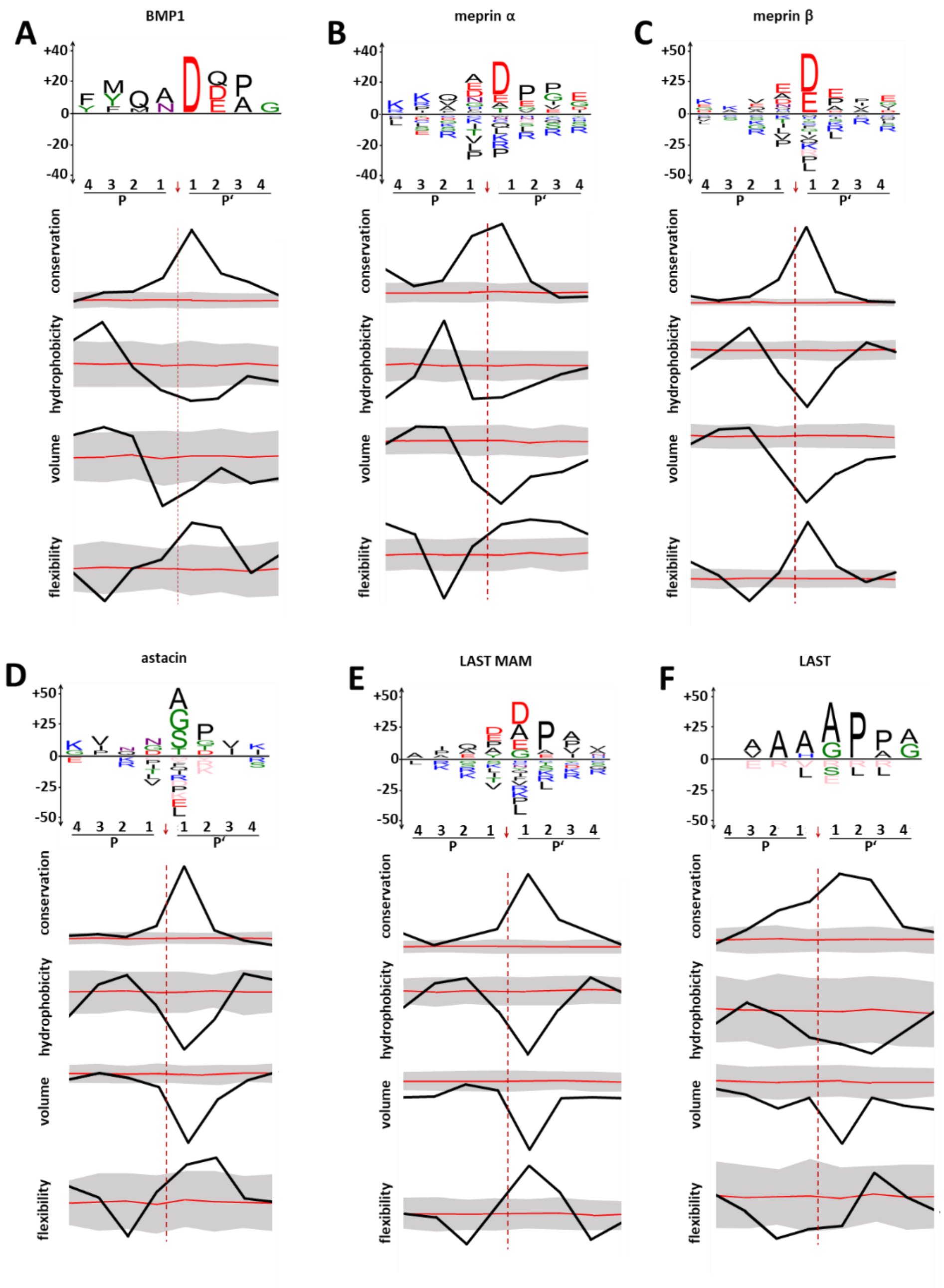


**Supplemental Figure 5. Physicochemical properties of astacin proteases.** Overview of the common cleavage preferences and physicochemical properties of BMP1 (A), meprin α (B), meprin β (C), astacin (D), LAST MAM (E) and LAST (F). The analysis was performed based on of the cleavage sites deposited in the MEROPS database (BMP1 (P13497), n=25; meprin α (Q16819), n=700; meprin ß (Q16820), n=879, astacin (P07584), n=199, LAST_MAM (B4F320), n=415; LAST (B4F319), n=76). Depicted in each case is a sequence logo (upper panel) displaying the difference to the natural abundance of residues in the cleavage site positions P4-P4', the degree of conservation of positions according to the BLOSUM62 substitution matrix (43) (upper graph), the hydrophobicity of the residues (44) (second graph from the top), the volume of the residues (45) (second graph from the bottom), and the flexibility of the residues (46) (bottom graph). The significance level is 95%. The error range is displayed in gray. The diagrams allow a semi-quantitative evaluation, since iceLogo does not allow scale normalization.

**A**

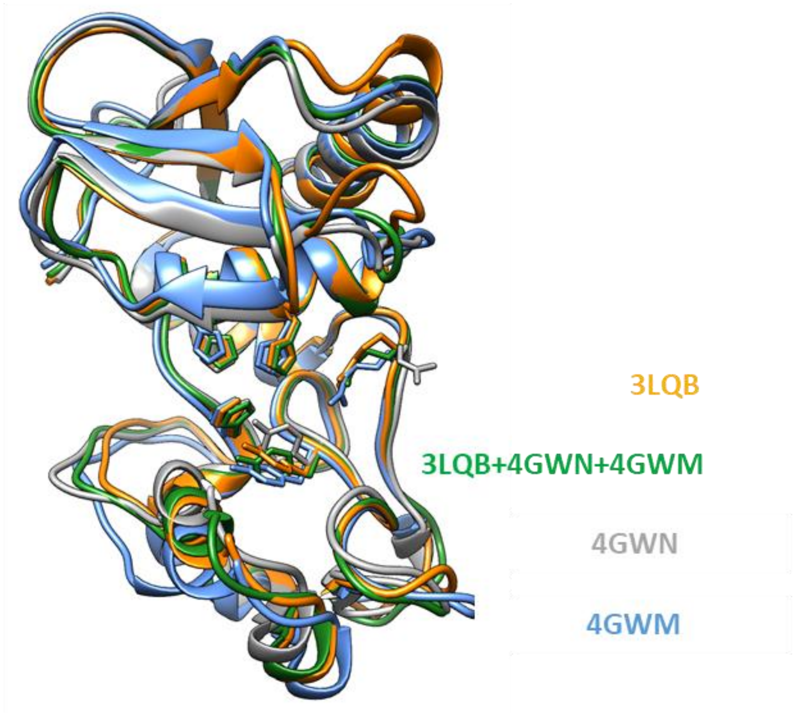


**B**

| BDP / residues in | 3LQB | 4GWN | 4GWM | 3LQB 4GWN 4GWM |
|---|---|---|---|---|
| strongly favored regions | 93.9 % | 89.1 % | 91.5 % | 92.7 % |
| favored regions | 4.8 % | 8.5 % | 7.9 % | 6.7 % |
| allowed regions | 0 % | 1.8 % | 0 % | 0.6 % |
| not allowed regions | 1.2 % | 0.6 % | 0.6 % | 0 % |

**Supplemental Figure 6. Structure modelling.** (A) Structural overlay of ovastacin homology models generated with template 3LQB (ZHE-1) is depicted in orange, with template 4GWN (meprin ß without propeptide) in gray, with template 4GWM (meprin ß with propeptide) in blue, and from the combined templates 3LQB, 4GWN, and 4GWM in green. Side chains of histidine 182, 186, and 192, as well as tyrosine 238 and arginine 264 are displayed. (B) Quality check of models generated in (A) with PROCHECK (49, 50) calculating the fraction of residues of a model with permissible torsion angles in the Ramachandran plot and thus allows an assessment of the plausibility of a model. If more than 90% of the residuals are in the strongly preferred range, a good model quality can be assumed.
